# Characterization of Human Anterior Neural Organoids as a Model for Investigating Cohen Syndrome

**DOI:** 10.1101/2025.04.23.650357

**Authors:** Renuka Prasad, Ju-Hyun Lee, Da-Yeon Lee, Yeon-hee Lee, Jihae Lee, Si-Hyung Park, Jae Ryun Ryu, Boram Lee, Seungji Choi, Jungmin Choi, Il-Joo Cho, Joon-Yong An, Fabrizio Vacca, Muhammad Ansar, Hyun Jung Kim, Myungjin Kim, Woong Sun

**Author notes:** Equally contributed as co-first authors (Renuka Prasad, Ju-Hyun Lee). Correspondence to: Woong Sun, Ph. D. Department of Anatomy, Korea University College of Medicine, Seoul, 02841, Republic of Korea, Correspondence may also be addressed to: Myungjin Kim, Ph.D Korea Brain Research Institute, Daegu, 41068, Republic of Korea.

## Abstract

Neural organoids display three-dimensional (3D) structures that resemble in vivo neural architectures. Previously, we developed a novel two-dimensional (2D) neural induction-based protocol for culturing spinal cord organoids, enabling size control and recapitulating neural tube morphogenesis. In this study, we evaluated the application of this concept to induce the anterior regions of the brain and generate human anterior neural organoids (hANOs). By inducing neuroepithelial (NE) cells in 2D and re-aggregating them led to tube-forming morphogenesis similar to that posterior spinal cord induction. The transcriptome profiles of these hANOs resembled the frontal cortex of 20 weeks post-conception (PCW) human embryos. Using this hANOs protocol, we investigated microcephaly phenotypes associated with Cohen syndrome (CS), caused by biallelic loss-of-function variants in *VPS13B* gene. Deleting VPS13B in human pluripotent stem cells resulted in Golgi dispersion and growth retardation onset in mutant hANOs, akin to CS patients with postnatal microcephaly. This delay is partly linked to reduced neuronal growth. Additionally, mature CS organoids showed enhanced hyper-excitability associated with an excitatory/inhibitory imbalance. In conclusion, this protocol is suitable for studying microcephaly phenotypes from human genetic mutations due to its simplicity and scalability.

## INTRODUCTION

Cohen syndrome (CS) is a rare autosomal recessive disorder characterized by intellectual deficiency, postnatal microcephaly, dysmorphic facial features, hypotonia, neutropenia, and progressive retinopathy.^1^ CS is caused by biallelic loss of function (LOF) mutations of the *VPS13B* gene. ^2,3^ VPS13B belongs to the Bridge-Like Lipid Transfer Proteins (BLTPs) family, which is proposed to mediate lipid transport at intracellular membrane-contact sites^4^ The VPS13B plays an essential role in the assembly and maintenance of the Golgi complex.^3,5^ Specifically, the VPS13B protein is implicated in the glycosylation, as well as sorting, and transportation of lipids and proteins within the cell.^6^ Consequently, cells from CS patients display the characteristic cellular phenotype of Golgi dispersion.^5^ However, the precise mechanism by which VPS13B deficiency or these cellular changes give rise to the clinical manifestations of CS remains unclear.^7^ Although modeling VPS13B knockout (KO) rodent models have provided valuable insights into its roles in the mammalian brain, the postnatal microcephaly observed in CS patients is either mild or not replicated in mice.^8–11^ This limitation underscores the necessity for more faithful experimental models to advance our understanding of CS pathology.

Microcephaly is a clinical condition characterized by reduced head and brain size, particularly the frontal cortex are smaller than in a typically developing child.^12^ Microcephaly is classified into primary and secondary categories. Primary microcephaly is a congenital neurodevelopmental defect.^13^ Secondary microcephaly occurs during the postnatal period, and often results from maturation deficits. A wide array of factors contributes to microcephaly, including toxic exposure to alcohol, in utero infections (notably Zika virus), and genetic or metabolic issues.^13–15^ Brain organoids derived from induced pluripotent stem cells (iPSCs) of patients harboring mutations in critical genes, such as *CDK5RAP2, ASPM, WDR62*, and *CPAP*, offer a platform for elucidating the molecular mechanisms underlying primary microcephaly. ^16–19^ Secondary microcephaly, on the other hand, is often is associated with a group of disorders known as developmental and epileptic encephalopathies (DEE), which are linked to several genes.^20^ Cerebral organoids derived from DEE patient iPSCs exhibit impaired expression of cortical layer markers and layering defects. Additionally, dorsal forebrain organoids generated from iPSCs of a Rett syndrome patient with secondary microcephaly also showed premature development of the deep cortical layer.^21^ Overall, the brain organoid models faithfully recapitulate the pathological hallmarks of microcephaly within the context of human tissue and provide a deeper insight into microcephaly.

Recent advances in organoid culture protocols have enabled the generation of brain organoids to address important research questions such as the mechanism of microcephaly.^16,22–24^ The main limitation of current organoid production protocols is the high heterogeneity and the limited number of organoids produced per batch, both of which pose challenges for quantitative modeling of disorders.^25,26^ To overcome these challenges, we recently developed a protocol for generating human spinal cord organoids (hSCOs) using an initial induction of human pluripotent stem cells (hPSCs) in a 2D culture format rather than the traditional 3D embryoid body formation.^27^ Based on this protocol, hPSCs were exposed to dual SMAD inhibition, and a defined number of neurally induced cells were used for reaggregation. Notably, using this modification, we generated large numbers of uniformly sized human anterior neural organoids (hANOs), providing a reliable means of generating neural organoids of consistent quality and quantity suitable for microcephaly modeling.

Thus, we have characterized and utilized an improved hANOs culture system to model the pathology of CS and to address the mechanism of human secondary microcephaly caused by biallelic LOF mutations in the *VPS13B* gene.

## MATERIALS AND METHODS

### Target sgRNAs cloning into Cas9 plasmid

The sgRNA pairs were carefully designed using online CRISPR tools (CHOPCHOP and CRISPOR) to specifically target VPS13B gene while predicting any potential off-target effects.^28^ Exon 4 was the target region. The two sgRNA sequences were about 85bp apart. The complementary oligonucleotide pairs for sgRNA #1 and #2 were annealed at 95°C for 5 min, with ramp-down to 25°C to form a dsDNA fragment, which was ligated into AgeI, XbaI-digested lentiCRISPR v2 (Plasmid #52961, Addgene).^29,30^

### Human PSC culture and transfection

The H9 hPSCs were cultured on Matrigel-coated plates (Corning, 354277) in mTeSR1 medium (STEMCELL Technologies, 85850). Cells were maintained at 37°C in 5% CO2, with daily medium changes. The cells were passaged every 4 to 6 days using ReLeSR (STEMCELL Technologies, 05872), replating small cell clusters onto Matrigel-coated dishes. Cells at 60% confluency were transfected with the gRNA vector using Lipofectamine Stem Transfection Reagent (Invitrogen) according to the manufacturer’s protocol. The cells were passaged with Accutase (STEMCELL Technologies, 07922) and plated as single cells in the presence of 10 µM Rock inhibitor (Tocris, 1254). Three days later, the medium was replaced with hPSCs medium containing 1 μg/mL puromycin. After 48 hours of selection, daily medium changes continued until the colonies were large enough to passaged. Following puromycin selection, colonies were picked and expanded in six-well plates. Screening and characterization of the clones were performed based on their morphology and pluripotency marker expression patterns.

### Screening for deletion mutants and off-target analysis of hPSCs line

Genomic DNA (gDNA) was extracted from transfected hPSCs with the AccuPrep Genomic DNA Extraction Kit (Daejeon) according to the manufacturer’s instructions. Genome deletions resulting from the sgRNA pairs were detected by AccuCRISPR In/del analysis service (Bioneer). The PCR primers targeting the different exons of the VPS13B transcripts are used for confirmation of the gene KO (Supplementary Table 1). Potential off-target sites were selected with CRISPOR online tools. The sites were PCR-amplified prior to Sanger sequencing. The off-target sites with three mismatches compared to each sgRNA were sequenced.

### Generation of human anterior neural organoids

Human anterior neural organoids (hANOs) were generated according to a previously published protocol with slight modifications.^27^ Briefly, hPSCs were passaged by ReLeSR into small clumps and replated onto matrigel-coated plates in mTeSR1. After 2 days, mTeSR1 was replaced with differentiation medium (DM), consisting of DMEM/F-12 (Life Technologies, 11320033), 1% N2 (Life Technologies, 17502048), 2% B27 (Life Technologies, 17504044), 1% nonessential amino acids (NEAA) (Life Technologies, 11140050), 1% penicillin/streptomycin (P/S) (Life Technologies, 15140122), and 0.1% β-mercaptoethanol (Life Technologies, 21985023). To differentiate into anterior neuronal lineages, cells were treated with SB431542 (10 uM, TOCRIS, 1614) and LDN193189 (0.1 uM, Stemgent, 04-0074) in DM for 3 days with daily media change. On day 3, intact colonies were gently detached by 2.4 unit/ml Dispase II (Life Technologies, 17105041) treatment for 20 min at 37□. Detached colonies were then transferred onto uncoated culture dishes in DM supplemented with basic fibroblast growth factor (bFGF) (20 ng/ml, R&D Systems, 233-FB) and 20ng/mL epidermal growth factor (EGF; Gibco, PHG0311). To generate VPS13B KO hANOs, cell clumps were dissociated into single cells using Accutase after 3 days of neural induction. A total of 5,000 dissociated cells were seeded into each well of a 96-well low-attachment plate supplemented with bFGF and FGF. The hANOs were fed daily for 4 days. On day 7, hANOs were cultured in DM without bFGF and EGF. After 8 days, hANOs were transferred to a 60-mm petri dish onto an orbital shaker and grown in 1:1 mixture of DMEM/F-12 and neurobasal medium (Life Technologies, 21103-049; containing 0.5% N2, 1% B27, 0.5% NEAA, 1% P/S, 0.1% β-mercaptoethanol and 1% GlutaMAX (Life Technologies, 35050-061). Media was changed every 3 to 5 days. We followed the guideline for the production of neural organoids.^31^

### Reverse Transcriptase-PCR

Examination of gene expression was carried out by RT-PCR. Total RNA was isolated using TRIzol reagent (Invitrogen) according to the manufacturer’s instructions. cDNA was synthesized from 1µg RNA using Moloney murine leukemia virus reverse transcriptase (MMLV, Promega, Madison, WI) with random primers. cDNA was amplified using gene-specific primers (Supplementary Table 2).

### Primary neuron culture from organoids

For the primary human neuron culture, cells were dissociated from around 1-month-old organoids using Cysteine-papain-solution (Worthington) and plated onto Poly-L-Ornithine/laminin coated coverslips at a density of 100,000 cells/well in 24-well plate in base media consisting of Neurobasal media, GlutaMAX medium (Thermo Fisher) supplemented with B27 supplement (Thermo Fisher), 1% penicillin/streptomycin (P/S). Next day, cells are fed with base media containing ascorbic acid phosphate, cAMP, Laminin (Invitrogen), Culture one (Invitrogen), Chemically Defined Lipid Concentrate (Invitrogen), Thereafter, one-half of the medium volume was changed every 3–4 days. Neurite length was measured using the semi-automated tracing tool, NeuronJ (a plug-in for ImageJ).

### Immunofluorescence staining

The hANOs were fixed by immersion in 4% paraformaldehyde (PFA, Biosesang) overnight at 4□ and washed several times in PBS. The fixed organoids were then incubated in 30% sucrose in PBS at 4□ until completely submersed, embedded in Tissue-Tek Optimal Cutting Temperature (O.C.T. Compound, SAKURA), frozen on dry ice, and cryosectioned serially at 16- to 40-µm thickness and collected onto New Silane III coating slides (Muto Pure Chemicals Co. Ltd, 5118-20F). For immunostaining, samples were permeabilized with PBST (0.1% Triton X-100 in PBS) three times for 5 min each at RT, blocked with solution (3% BSA, 0.2% Triton X-100, 0.01% sodium azide in PBS) for 30 min at RT, and then incubated with the respective primary antibody diluted in blocking solution for overnight at 4□ (Supplementary Table 3). Samples were then washed with PBST three times for 5 min each at RT, and then incubated with the respective secondary antibody and Hoechst33342 diluted in blocking solution for 30 min at RT. The secondary antibody was subsequently washed with PBST, and the samples mounted in Crystal mount (Biomeda, M02). All steps were performed with gentle shaking. Images were taken and processed using a Leica TCS SP8 Confocal microscope system.

### 3D whole-mount imaging and clearing

The organoids were fixed with 4% PFA in PBS for 30 min (<500 µm diameter) or 1 hr (>500 µm), then washed several times with PBST, and incubated with blocking solution (6% BSA, 0.2% Triton X-100, 0.01% sodium azide in PBS) for overnight. For 3D wholemount immunostaining, samples were immersed in primary antibody diluted in blocking solution for 48 hrs. The primary antibody was then washed with PBST three times for 10 min each. The samples were then incubated with the appropriate secondary antibody and Hoechst33342 diluted in blocking solution for 48 hrs. Subsequently, samples were washed with PBST three times for 10 min each and mounted onto cover glass (24 x 40 mm) with mounting solution (25% urea and 65% sucrose in H_2_O) for optical clearing. All steps were performed in a 0.2 ml PCR tube with gentle shaking at RT. All images were captured with a Leica TCS SP8 Confocal microscope.

### 3D Image processing

For 3D whole imaging, raw images were collected using a Leica SP8 and processed with LAS X software (Leica). The organoid images were created from Z-stacks (typically 50–300 images) with 0.5–2 µm intervals, and then manually segmented and rendered in Leica software. The regions of interest (ROI) were manually segmented and visualized using 3D reconstructions in Leica.

### Bulk RNA-seq analysis

Raw RNA-Seq data were processed using Salmon v1.10.0 to generate count matrices against the human reference genome (GRCh38/hg38).^32^ The resulting count matrices were normalized and preprocessed using the regularized log (rlog) transformation implemented in DESeq2 v1.38.3. ^33^ To mitigate batch effects and remove unwanted technical variability, we employed a combination of ComBat-seq, surrogate variable analysis (SVA), and Limma.^34–36^ Differential gene expression analysis was conducted using DESeq2. Genes with an absolute log2 fold change ≥ 0.5 and adjusted p-values < 0.05 (Benjamini-Hochberg correction) were considered differentially expressed. Results were visualized using EnhancedVolcano v1.16.0. Hierarchical clustering of DEGs was performed using DEGreport v1.39.6. Gene set enrichment analyses (GSEA) were performed using fgsea v1.27.0 on the Gene Ontology Biological Process gene set. An adjusted p-value threshold of 0.05 was used to determine significant enrichment. enrichMate algorithm was implemented to visualize the GSEA results. Heatmaps were generated using the pheatmap v1.0.12 to visualize expression patterns.^37^

For brain similarity analysis, we obtained normalized microarray data from the Allen Brain Institute (human.brain-map.org). The datasets cover 27 brain regions, which we grouped into 10 major regions, including the prefrontal, frontal, parietal, temporal, and occipital cortices, as well as amygdala, hippocampus, striatum, thalamus, and cerebellum. RNA profiles from neural organoids were compared with the microarray dataset using Spearman Rank Correlation Coefficients. Temporal similarity was assessed by comparing the expression profiles of the human frontal cortex across different post-conception weeks.

### Single-nuclear RNAseq data analysis

Nuclei were isolated from triturated organoids cell suspension in NbActiv1 (Brainbits LLC, Illinois). Briefly, cells were spun down at 500 rcf for 5 min at 4°C. The cell pellet was mixed with chilled 0.1X Lysis Buffer (1X LB: 10 mM Tris-HCl (pH 7.4), 10 mM NaCl, 3 mM MgCl2, 0.1% Tween-20, 0.1% NP-40, 0.01% Digitonin, 1% BSA in Nuclease-free Water) and incubated for 5 min on ice. After adding Wash buffer (10 mM Tris-HCl (pH 7.4), 10 mM NaCl, 3 mM MgCl2, 0.1% Tween-20, 1% BSA in Nuclease-free Water), cells were mixed and spun down at 500 rcf for 5 min at 4°C. The cell pellet was recovered and washed twice more for a total of 3 washes. Cells were resuspended in a chilled Diluted Nuclei Buffer (10x Genomics) and passed through a 40 µm Flowmi Cell Strainer (Bel-Art). The nuclei concentration was determined using a Countess II FL Automated Cell Counter. Nuclei were visually inspected and counted using both the Ethidium Homodimer-1 fluorescent dye (LIVE/DEAD Viability/Cytotoxicity Kit, Thermo Fisher Scientific) and Trypan Blue stain (Thermo Fisher Scientific) according to the manufacturer’s instructions. Ten thousand single nuclei were immediately processed into the 10x Genomics Chromium Controller. Libraries were generated following the Chromium Next GEM Single Cell 3’ Gene Expression User Guide (v3.1 Chemistry Dual Index, 10x Genomics), analyzed using the 4200 TapeStation (Agilent), and sequenced on an Illumina HiSeq X.

Sequenced reads of sample hCOS01G were aligned to the GRCh38 2024-A human reference genome using Cell Ranger version 8.0.1 (10x Genomics) with default parameters. Cell Ranger detected a fraction of 93.1% reads and a median of 432 genes per cell. For quality control, we included genes expressed in at least 10 cells and cells expressing between 100 and 1,500 genes with a unique molecular identifier (UMI) count ranging from 200 to 2,000. Outlier cells with a proportion of mitochondrial genes over 0.4% and putative doublets were filtered by Scrublet (v0.2.1).^38^ Normalization and log transformation were performed using Scanpy (v1.9.8).^39^ PCA was applied, reducing the dimensionality of the data using 50 components. The nearest neighbor graph was computed, and the UMAP embedding was generated. Clustering was performed using the Leiden algorithm with a resolution of 0.8. The imputation method MAGIC was applied to improve gene expression representation.^40^ Cell types were identified based on the imputed expression profiles of known marker genes. SOX2, ASCL1, NEUROD6 for neural progenitor cells (NPC); SLC17A7 for excitatory neurons; GAD1, GAD2 for inhibitory neurons; PDGFRA, OLIG2, SLC17A3 for glial progenitor cells (GPC); and GFAP, AQP4, ALDH1A1 for astrocytes. Comparison of the transcriptional profiles of the organoid and the developmental human brain atlas was done by calculating Spearman Rank Correlation Coefficients between mean expression of genes within the clusters and the 11 stages of the human brain atlas.^41^ We then used Wilcoxon’s rank-sum test to assess if the stage of interest has significantly higher Spearman Rank Correlation Coefficients with the organoid clusters than the other stages. Only the genes expressed in all stages and clusters were used to calculate the correlations, and Benjamini-Hochberg (FDR) correction was applied for multiple comparisons. A gene set enrichment test was performed to compare the cell types of hCOS01G with the human brain atlas clusters. The background genes for the enrichment test were set to include all the overlapping genes of the human brain atlas and hCOS01G. This enrichment test included cluster-specific differentially expressed genes (DEGs) of the atlas and cell type-specific genes of hCOS01G, both attained with Wilcoxon rank-sum test in Scanpy, with a threshold of FDR <0.05 and >25% of cells within the cluster expressing the gene. A one-sided Fisher’s exact test with multiple comparisons by Bonferroni correction was applied.

### Electrophysiology

To investigate the functional differences between 5-month-old control organoids and VPS13B organoids, we used a MEMS neural probe integrated with 16 black Pt-coated microelectrodes to record neural signals.^42^ Each organoid was transferred from the culture dish into a 35-mm Petri dish-based recording chamber, where it was positioned beneath the neural probe. Once the organoid was in place, the neural probe was gradually inserted into the organoid using a microdrive.^43^ After insertion, the organoid and neural probe were immobilized in low-melt agarose gel, and 1.5 mL of fresh DMEM/F12-based culture medium was added to the chamber. The entire setup, including the organoid, was then enclosed in a small acrylic box to prevent rapid evaporation of the culture medium. All recordings were conducted in a CO□ incubator maintained at 37°C and 5% CO□.

Neural signals were recorded from the 16 black Pt-coated microelectrodes and amplified through an RHD2132 board connected to an RHD2000 Recording System (Intan Technologies, USA) with the following settings: 20 kS/s per channel, a 0.1 Hz high-pass filter and 300 Hz low-pass filter for local field potential (LFP) recordings, and a 300 Hz high-pass filter and 6 kHz low-pass filter for spike recordings. The system also utilized a 16-bit ADC. Neuronal activity from all organoids was recorded for at least 10 minutes. The recorded neural signals were analyzed to produce raster plots and detect single spikes using a custom MATLAB sorting algorithm developed previously. ^44^ A threshold amplitude of 50 μV, set at more than three times the noise level (∼10 μV), was used for spike detection.

Additionally, we analyzed raw local field potentials, amplitude spectrograms, filtered LFP signals, and LFP power spectrograms using a newly developed custom MATLAB sorting algorithm. Each signal was quantified and represented in a bar plot. Burst activity was evaluated with the ISIN-threshold method, using a 0.1 s ISI threshold and a minimum of three spikes per burst. All statistical analyses were verified with Student’s t-test using GraphPad Prism.

### Statistics

Statistical analyses were performed using an unpaired Student’s *t*-test to compare two groups, and with a one-way ANOVA with Tukey’s multiple comparison test for multiple comparisons. All analyses were performed using GraphPad Prism 8 software version 8.0.2 (GraphPad, CA, USA), and the results are presented as the mean□±□SEM or SD.

## RESULTS

### Anterior neural organoids exhibit tube morphogenesis with distinct morphological features

Our research team recently developed a new culture protocol for modeling hSCOs, through which we were able to create a tissue analogue of the posterior nervous system.^27,45^ This process was found to mimic the posterior nervous system, along with the induction of neuromesodermal progenitors. Considering that actual neural tube (NT) formation is a critical process in the development of the anterior nervous system, it is necessary to analyze in detail whether morphogenesis similar to NT formation occurs in models induced into the anterior nervous system through similar culture methods. Thus, we sought to examine whether anteriorly derived spheroids also undergo NT morphogenesis. To mimic organoids resembling the anterior nervous system, we subjected hPSCs monolayers to dual SMAD inhibition, inducing neuroepithelial (NE) cell formation in 2D (Fig. 1a). The NE cells were then detached using dispase and aggregated with FGF2 and EGF to form reaggregates. In these reaggregates, NE cells self-organize and proliferate into 3D structures with follicle-like morphology within anterior spheroids. These spheroids subsequently differentiate and mature into hANOs.

**Fig. 1.**
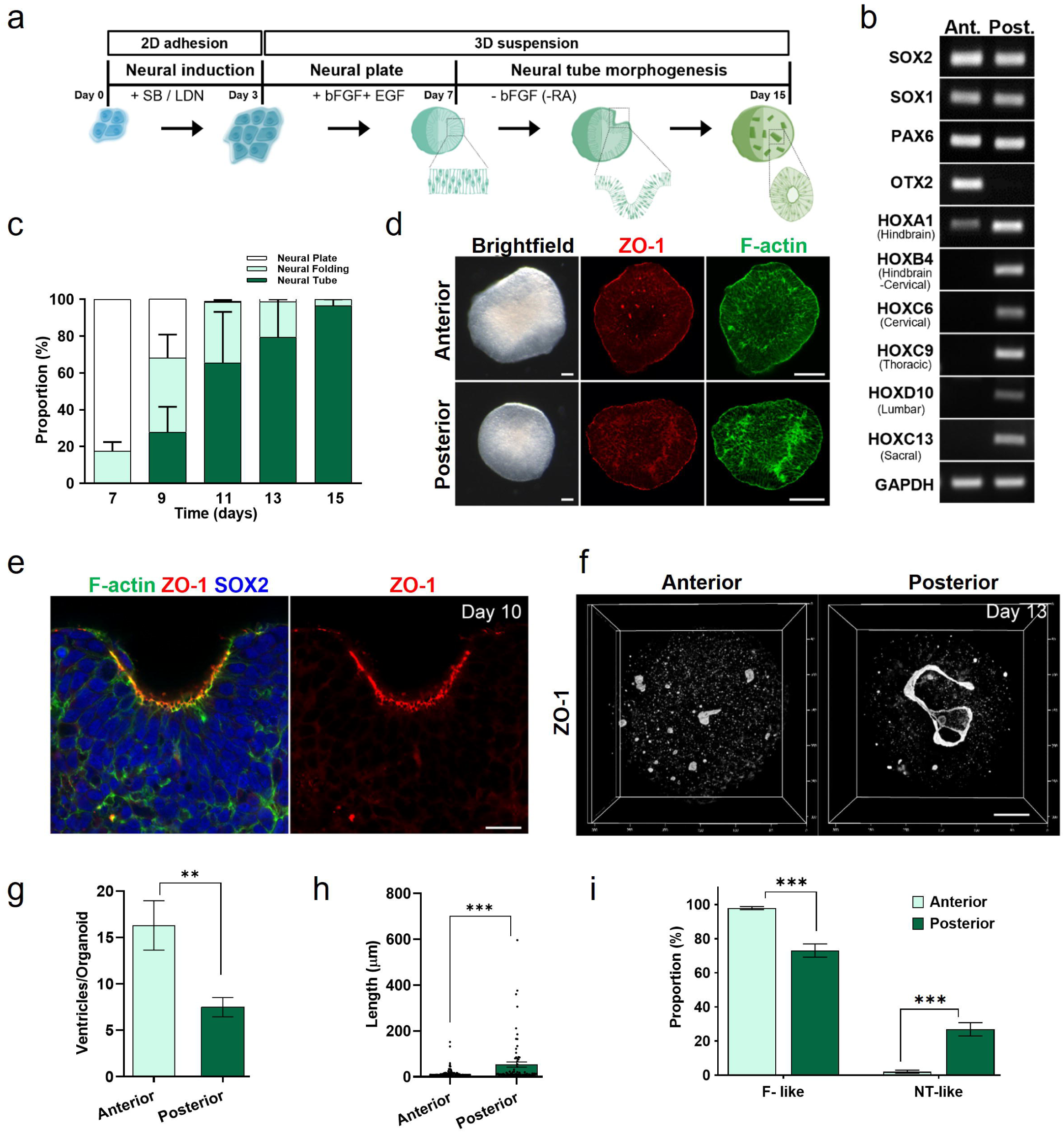
Comparison of anterior with posterior organoids. **a** Schematic of the procedure for generating anterior organoids. Instead of using SB+CHIR for posterior NSC induction, we used SB+LDN to induce anterior NSCs. **b** Expression of regional markers of anterior versus posterior NSCs from hPSCs. **c** Quantification of the proportion of anterior organoids at different stages at indicated days of culture. **d** Comparison of anterior with posterior organoids at the NP stage. Polarization of the surface of the NE layer was visualized via staining for F-actin (green) and ZO1 (red). Scale bar, 100µm. **e** Neural fold of anterior organoids. The NE was visualized with SOX2 (blue), F-actin (green), and ZO1 (red) staining. Scale bar, 20µm. **f** 3D morphology of NTs labeled with ZO1 staining in anterior versus posterior organoids. Scale bar, 50µm. **g** Quantification of the number of ventricles that include follicle-like and neural tube–like structures in organoids. (Unpaired t-test; *p* < 0.01, n=10). **h** Scatter dot plot of individual ventricle length. (Unpaired t-test; *p* <0.001, Rostral n=163, Posterior n=75). **i** The proportion of follicle-like and neural tube–like ventricles in the anterior versus posterior organoids (Unpaired t-test; F-like, *p* <0.001, n=10; NT-like, *p*<0.001, n=10).

Both protocols produced NPC expressing pan-NPC markers such as SOX2, SOX1, and PAX6. However, anterior NSCs selectively expressed the anterior marker OTX2 (Fig. 1b). In our protocol, anterior spheroids also underwent NT morphogenesis (Fig. 1c), exhibiting proper NE polarization (Fig. 1d) and neural fold formation (Fig. 1e). Interestingly, the resultant hANOs from anterior induction protocols exhibited distinct ventricular morphology compared to hSCOs from posterior induction. While posterior hSCOs exhibited long and connected NT-like morphology, anterior hANOs showed short, isolated, follicle-like morphology (Fig. 1f-i). The mechanisms underlying these differences remain unclear. However, these findings suggest that the neurulation process is region specific, and that self-organizing properties of regional stem cells mediate spinal cord elongation and the morphogenesis of the follicular brain hemisphere.

### hANOs display cellular diversity and synaptic maturation

We next examined whether the hANOs generated using our protocol contained typical cell types found in cerebral organoids. At 1-month, the NPC marker SOX2 was expressed in the proliferative NE (Fig. 2a). By 4-months, the organoids revealed a significant population of astrocytes, indicated by GFAP expression, along with differentiated neurons. Additionally, the axons of the differentiated neurons also labeled with the neurofilament M (NF-M) (Fig. 2b). The inhibitory neuron marker GABA was also detected at this stage. We also performed an analysis of markers for neuronal subtypes within cortical plate (CP)-like regions. As the hANOs developed, the rosette structure became less prominent, and the laminar organization disappeared. However, CTIP2^+^ deep-layer neurons and SATB2^+^ superficial neurons were frequently clustered distinctly, reflecting remnants of the laminated features of the cerebral cortex (Fig. 2b). Moreover, mature synapses were observed in the hANOs, as indicated by closely associated puncta of the pre-synaptic marker SV2A and the post-synaptic marker PSD95. These puncta were well-aligned with the mature neuronal dendritic marker MAP2 (Fig. 2c), demonstrating neuronal synaptic maturation.

**Fig. 2.**
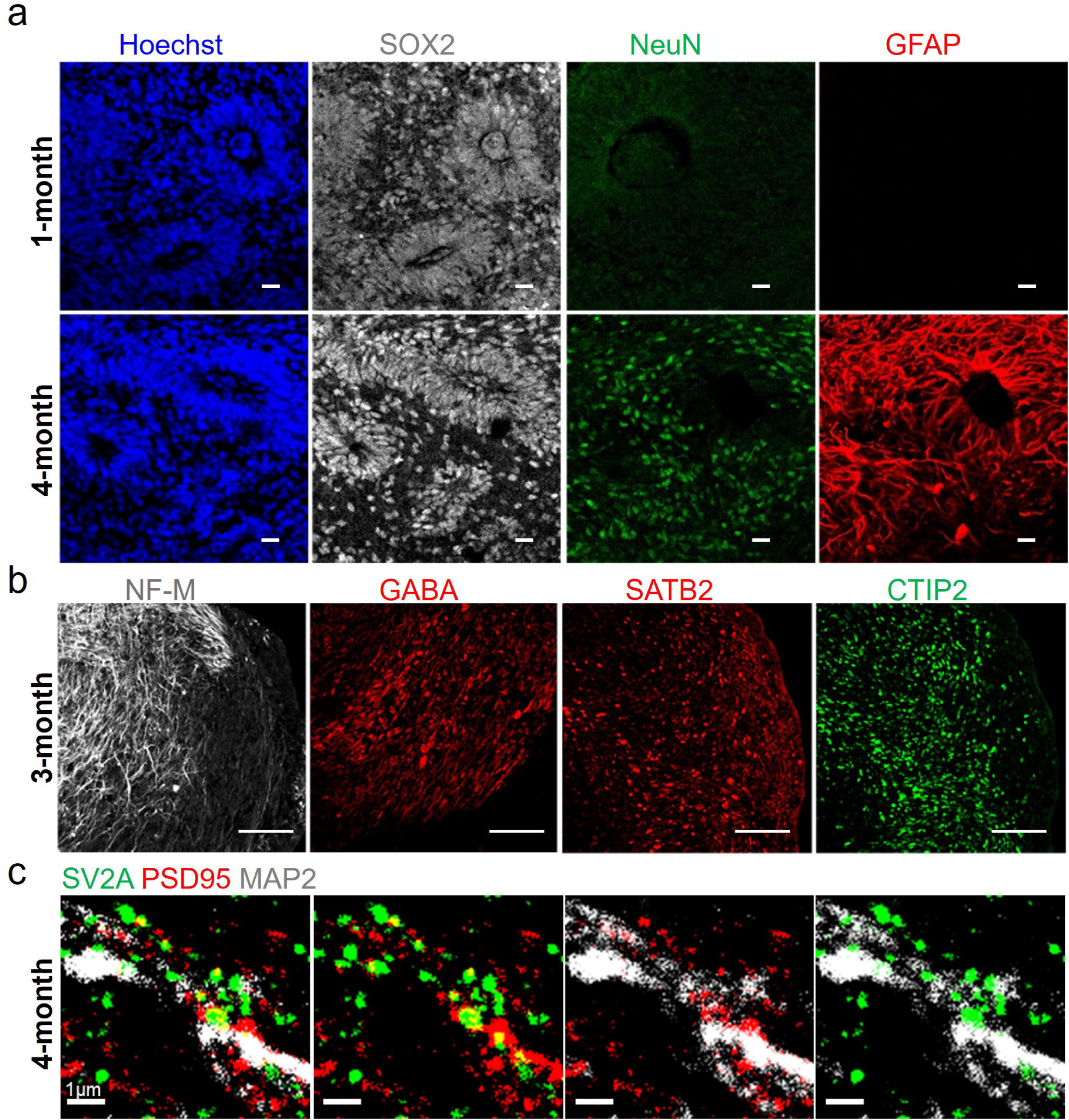
Validation of hANOs. **a** Immunostaining of 1-month organoid shows SOX2^+^ NPC lining the ventricle-like structures (shown by dashed lines) within hANOs. 4-month organoids show significant presence of GFAP^+^ astrocytes and the mature neuronal marker NEUN at this developmental stage. Scale bar, 20um. **b** Immunostaining of 3-month hANOs shows the presence of neuronal marker NF-M and inhibitory neuronal marker GABA. Scale bar, um. The 3-month hANOs also show the presence of the superficial SATB2^+^ cortical layer and deeper layer CTIP2^+^ zone. Scale bar, 75µm. **c** Immunostaining of 4-month hANOs for synaptic maturation show the expression of the pre-synaptic marker, synaptic vesicle (SV2A) and postsynaptic marker, PSD95 which is partially co-localized with mature dendritic marker MAP2. Scale bar, 1µm. (n = 3 independent organoids).

### Transcriptomic comparison of hANOs with human fetal brain regions

To characterize the hANOs in terms of brain similarity, we performed bulk RNA sequencing of neural organoids derived from both iPSCs or H9 cells at different time points. These results were compared with normalized microarray datasets from the Allen Brain Institute.^46^ Our analysis revealed that the transcriptional profiles of hANOs closely resemble those of cortical regions, while also exhibiting broad similarity to non-cortical areas in the developing human brain (Fig. 3a). Furthermore, we examined the transcriptional profiles of hANOs in relation to the temporal expression patterns of the frontal cortex from 19 to 21 weeks post-conception (PCW), which corresponds to the second trimester of human pregnancy (Fig. 3b), revealing a trend of increasing similarity over time.

**Fig. 3.**
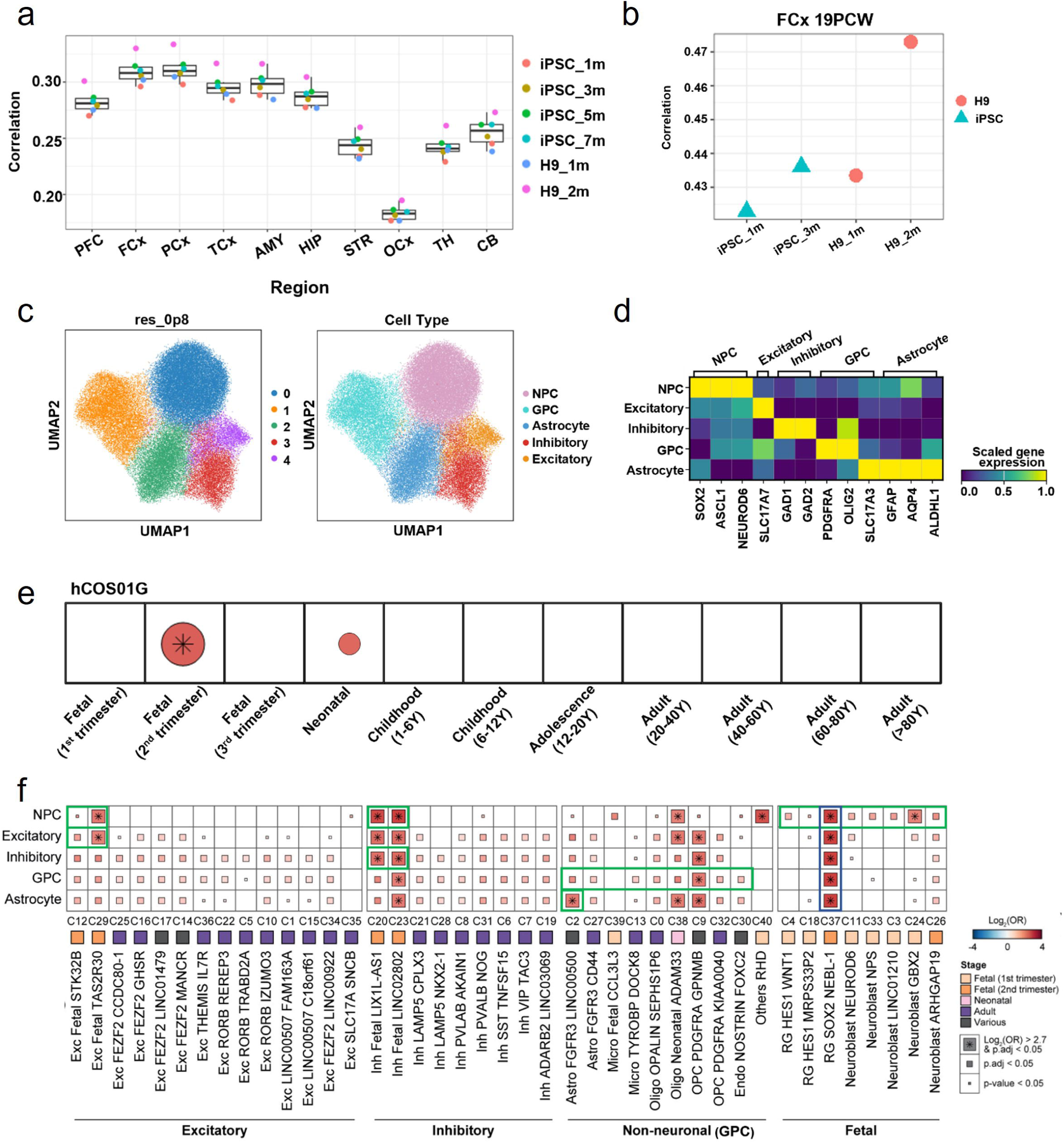
Transcriptomic comparison of hANOs with human fetal brain regions. **a** Predicted regional identity of ANOs based on correlation. **b** ANOs gene expression corresponded to the frontal cortex (FCx) of human embryo at 19-24 weeks post-conception (PCW). **c** Uniform manifold approximation and projection (UMAP) of single-nucleus RNA (snRNA) data from hCOS01G, colored by annotated cell types. **d** Scaled expression of marker genes across cell types. **e** Wilcoxon rank-sum test of Spearman correlations comparing gene expression of hCOS01G cell types with 11 developmental stages from the human brain atlas, with Benjamini-Hochberg (FDR) correction for multiple comparisons. **f** Gene set enrichment analysis of cell type markers from the atlas, using a one-sided Fisher’s exact test with Bonferroni correction for multiple comparisons.

To delineate the cell types in the hANOs, we performed single-nucleus RNA sequencing on cells from a 5-month-old organoid (hCOS01G), identifying five distinct cell types: NPC, excitatory neurons, inhibitory neurons, glial progenitor cells (GPC), and astrocytes (Fig. 3c). Each cell type showed distinct expression of well-established markers (Fig. 3d), which supports our experimental findings shown in figure 2 regarding the populations of progenitor cells, astrocytes, inhibitory neurons, and excitatory neurons in hANOs.

To assess the resemblance of the organoid with different developmental stages in the human brain, we compared its transcriptional profiles with those from the human brain atlas, which covers brain cell types across 11 developmental stages from 7 gestational weeks to 90 years age.^41^ We observed a strong similarity between the expression profiles of the organoid and the fetal second trimester of human development (correlation coefficient = 0.845, p.adj = 1.26 × 10^-5^) (Fig. 3e and Supplementary Table 4). Further analysis of cell type-specific enrichment between the organoid and human brain yielded consistent results, showing that organoid cell types are highly enriched with the fetal second trimester clusters of the atlas. All organoid cell types were significantly enriched with the atlas cluster C37, which represents the fetal second trimester radial glia cell population. NPC also showed significant overlap with neuroblasts (C24, Log_2_(OR) = 2.71, p.adj = 8.50 × 10^-7^) as well as with fetal second trimester neurons (C20, Log_2_(OR) = 3.87, p.adj = 1.41 × 10^-14^; C23, Log_2_(OR) = 4.13, p.adj = 1.12 × 10^-16^; C29, Log_2_(OR) = 2.91, p.adj = 1.43 × 10^-7^) in the atlas. Excitatory neurons in the organoid exhibited expression profiles highly similar to those of excitatory neurons in the fetal second trimester brain (C29, Log_2_(OR) = 2.71, p.adj = 1.16 × 10^-7^), while inhibitory neurons strongly matched with the atlas inhibitory neurons of the same stage (C20, Log_2_(OR) = 3.06, p.adj = 1.06 × 10^-36^; C23, Log_2_(OR) = 3.20, p.adj = 1.84 × 10^-39^). Non-neuronal cells, such as GPCs and astrocytes substantially aligned with the corresponding non-neuronal clusters in the atlas. The oligodendrocyte precursor cell (OPC) cluster, which emerges in various stages of the human brain, was enriched in the organoid GPC (C9, Log_2_(OR) = 2.83, p.adj = 2.90 × 10^-31^) and astrocytes (C9, Log_2_(OR) = 3.07, p.adj = 8.74 × 10^-33^). Astrocytes were also specifically enriched in the astrocyte cluster of the atlas (C2, Log_2_(OR) = 2.88, p.adj = 9.26 × 10^-33^) (Fig. 3f and Supplementary Table 5). These results demonstrate that the transcriptomic profile of the 5-month organoid closely aligns with that of the developing fetal brain during the second trimester. This validates its developmental resemblance to this critical period and supports our previous findings of its resemblance to the 20 PCW frontal cortex.

### VPS13B KO organoids showed microcephaly-like feature

To investigate whether the hANOs culture protocol can effectively model disease symptoms including microcephaly found in CS patients, we generated VPS13B KO in H9 hPSCs using the CRISPR/Cas9 tool (Supplementary Fig. 1). Previous studies have shown that VPS13B protein is essential for Golgi integrity, and its knockdown results in Golgi dispersion.^5^ The VPS13B KO hPSCs showed Golgi dispersion, confirming its ablation in the VPS13B KO hPSCs (Fig. 4a, b). To compare the growth of hANOs from VPS13B KO and control cells, equal numbers (∼5000, initial cell number) of dissociated single NEs were used to generate reaggregates, which is known to improve homology of the organoid sizes.^2^^7^ At the 1-month stage, both the control and VPS13B KO hANOs exhibited similar size and morphology. Interestingly, the 2-month stage, VPS13B KO organoids were smaller in size compared to controls (Fig. 4c, d). This size difference between the two genotypes consistently maintained over time. We next investigated the mechanism underlying the smaller size of VPS13B KO organoids. To validate whether the proliferation of neural stem cells was predominantly affected by VPS13B KO, we assessed hANOs neural rosettes at 1-month stage by immunofluorescence for Ki67, a marker of proliferation. We observed comparable populations of Ki67^+^ cells in rosettes from VPS13B KO organoid compared with control organoids (Fig. 4e and Supplementary Fig. 2a). Next, we examined cell death in the hANOs using cleaved caspase-3 staining to detect early-stage apoptosis. We also found cleaved Caspase-3-positive cells were unchanged in VPS13B KO organoid sections (Fig. 4f and Supplementary Fig. 2b). These results indicate that VPS13B is not essential for NPC proliferation or survival, and that these factors were not major contributors to the growth retardation observed in VPS13B KO organoids. We next questioned whether altered neuronal growth could contribute to the smaller organoid size in the VPS13B KO group. To explore this, we assessed soma size by measuring the area around the NeuN-stained nucleus. At 2-months, the soma size of VPS13B KO neurons was significantly smaller compared to control neurons (Fig. 4g, h). Furthermore, it was reported that reduction of VPS13B leads to defective neurite outgrowth in cultured primary hippocampal neurons.^3^ This prompted us to test whether the changes in neurite length were also reflected in neurons derived from VPS13B KO organoid. Our analyses revealed that VPS13B KO neurons have a decreased average neurite length compared with controls starting from day 3 onwards (Fig. 4i, j). Together, these findings indicate that neurons in VPS13B knockout organoids exhibit noticeable reductions in soma size and neurite length, which may contribute to their microcephaly-like phenotype.

**Fig. 4.**
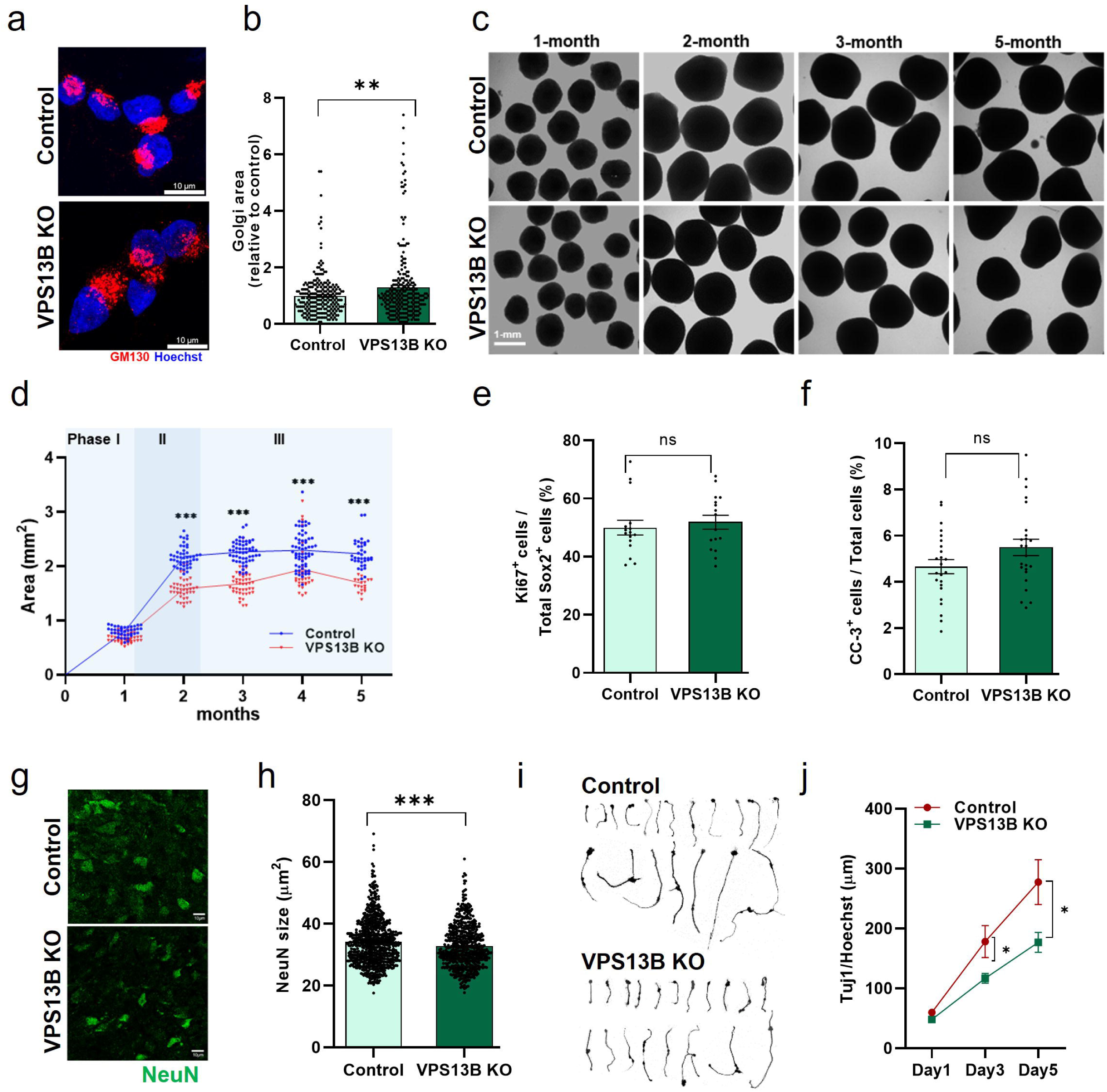
VPS13B KO hANOs show reduced size and impaired differentiation. **a** control and VPS13B KO hPSCs were stained with Golgi marker GM130 (red), and Hoechst (blue). Scale bars: 10 μm. **b** Quantification of Golgi areas of individual VPS13B KO cells relative to control cells (two-tailed unpaired *t*-test; \**p*□<□0.03, Golgi areas of n=240 cells were measured). **c** Representative images of control and VPS13B KO organoids at months 1, 2, 3 and 5. VPS13B KO organoids showed reduced size in comparison to control organoids from 2-month stage onwards. Scale bar: 1 mm. **d** Quantification of areas of control and VPS13B KO organoids. Depicted is the organoid area (mm^2^) from 1 to 5-month. (1 to 4-month (n= at least 30 organoids) and 5-month organoids (n= at least 19 organoids) were analyzed per condition; ^∗∗∗^*p* < 0.001 via one-way ANOVA. **e** Quantification of Ki67^+^ cells out of total SOX2^+^ cells in control and VPS13B KO organoids (1-month stage organoids, data are shown as mean ± SEM: n= 5; number of examined fields = 25). **f** Quantification of cleaved caspase-3 cells (CC3^+^) in control and VPS13B KO organoids at 1-month (two-tailed unpaired *t*-test; *non-significant*; n= 5; number of examined fields = 25). **g** The reduced soma size in 2-month VPS13B KO organoids was visualized with NeuN staining (Scale bar: 10 μm). **h** Quantification of soma size. Data is presented as mean ± SEM (two-tailed unpaired *t*-test; \*\*\**p*□<□0.001, n= 4, at least 600 NeuN labelled cells were quantified in each group from at least 21 images). **i** Representative examples of neurite tracing of day 3 neurons dissociated from 1-month organoids. **j** Quantification of neurite length of dissociated neurons at day 1, day 3, day 5 timepoints (unpaired Mann-Whitney test; day 3, \**p*□<0.017; day 5, \**p*□<0.021; n=7 images were quantified in each group per time point).

### RNAseq analysis of 2-month VPS13B KO organoids

To compare the transcriptional profiles of VPS13B KO organoids with controls, we performed bulk RNA sequencing on 2-month-old hANOs. After batch correction, principal component analysis (PCA) of the normalized expression data showed a separation between VPS13B KO and control organoids (Supplementary Fig. S3a), indicating distinct transcriptional profiles resulting from the VPS13B KO. Differential gene expression analysis using DESeq2 identified 79 differentially expressed genes (DEGs) in VPS13B KO samples, compared to 54 DEGs in control samples (|Log□ fold change| > 0.5, adjusted p-value < 0.05) (Fig. 5a).^33^ Hierarchical clustering of the 20 most significantly differentially regulated genes (ranked by statistics value) revealed distinct expression patterns in VPS13B KO organoids. The downregulated genes included critical neurodevelopmental regulators (*PAX6, MEIS1*) and synaptic plasticity genes (*SYNGAP1, NCAM2*), indicating disruption in both neuronal differentiation and synaptic connectivity pathways. The downregulation of genes such as *MTURN, RSPO2, and CITED2*, which are involved in cell growth and differentiation, may underlie the reduced growth observed in VPS13B KO organoids.^47–49^ Conversely, upregulated genes comprised cytoskeletal components (*TUBA1B, FLNA*) and synaptic proteins (*SHISA6*), suggesting compensatory mechanisms to maintain neuronal architecture and function (Supplementary Fig. S3b). GSEA analysis uncovered an enrichment of the term, regulation of membrane potential, in VPS13B KO organoids, primarily driven by genes critical for ion channel and transport processes, and synaptic function. To establish disease relevance, we compared our VPS13B KO organoid transcriptional profile with publicly available data from iPSC-derived neurons of CS patients.^50^ We observed an overlap7 genes (*ELN, PLEKHG5, PTGDS, MFAP2, TENM2, PAPLN, SSPN*) between the organoids and iPSC-derived excitatory neurons (Fig. 5b, c), with prominent expression in the disease conditions. The limited overlap of upregulated genes is likely attributable to the differences between the transcriptome profiles of neuronal populations and those of 2-month-old hANOs, which exhibit tissue-like cellular heterogeneity. GSEA analysis revealed that VPS13B KO organoids were significantly enriched in pathways related to neurogenesis and cell-cell adhesion, while control organoids showed enrichment in pathways associated with RNA processing and protein refolding (adjusted p-value < 0.05) consistent with the known role of VPS13B in intracellular protein trafficking (Fig. 5d, e and Supplementary Table 6).^11,51^ Our transcriptome analyses indicate that the dysfunction of the VPS13B protein in organoids provides new evidence that this impairment may contribute to disrupted neuronal differentiation and synaptic connectivity, potentially underlying the cognitive deficits observed in CS.

**Fig. 5.**
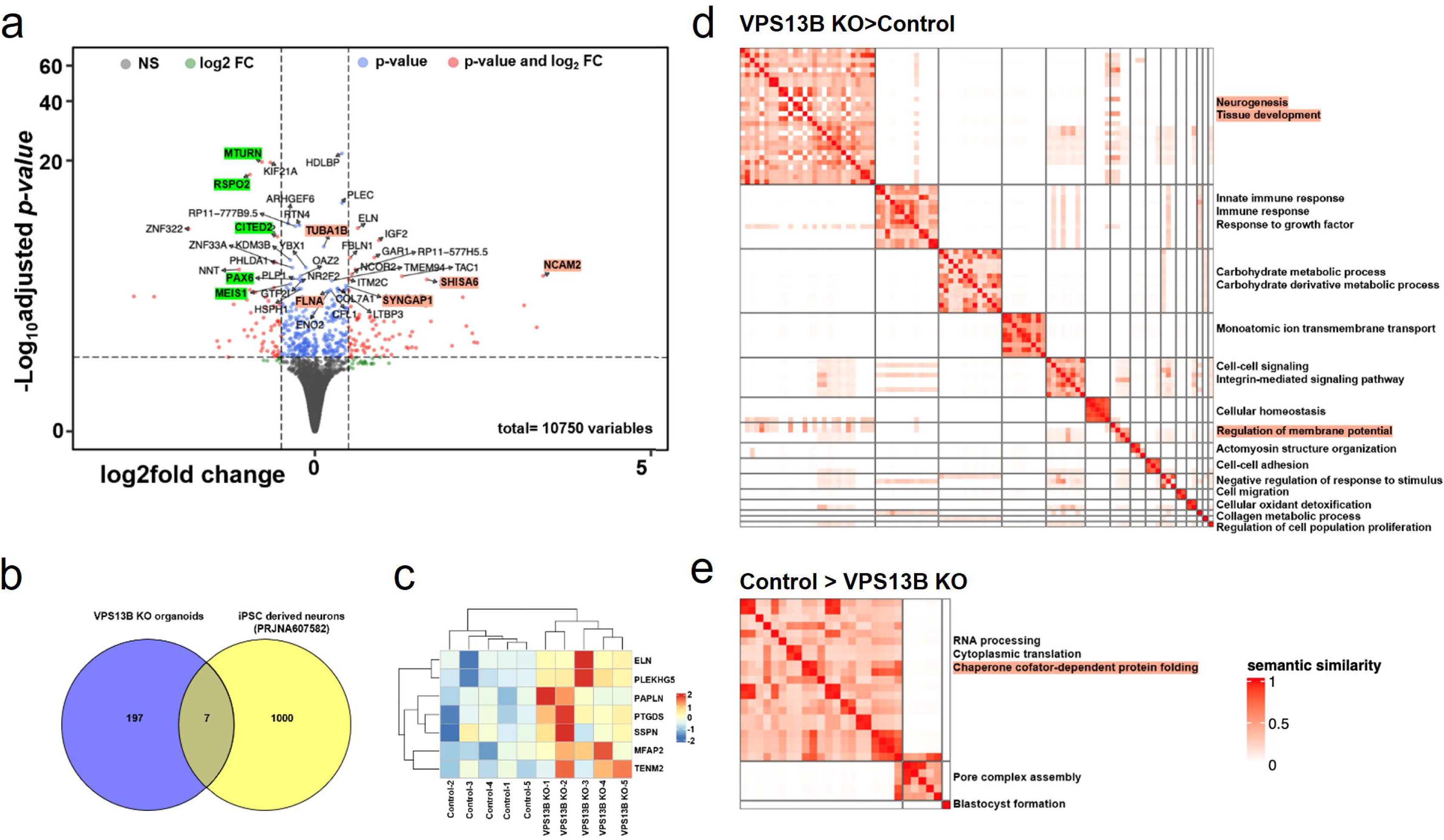
Differential gene expression and gene set enrichment analysis of control and VPS13B KO organoids. **a** Visualization of results from DGE test using volcano plot. X and Y axes show log2 fold change and −log10 adjusted p value, respectively. Left (Log2 fold change < 0) represents DEGs in control, and right (Log2 fold change > 0) represents VPS13B KO organoids. **b** Comparison of upregulated genes of the VPS13B KO organoids with publicly available iPSC-derived neurons from a patient with CS indicated 10 genes are commonly expressed. **c** Heatmap visualization of VPS13B KO organoid DEGs overlapping with iPSC-derived neurons from a CS patient. Gene expression intensities are displayed as colors ranging from blue for the lowest expression level to red for the highest expression level, as shown in the legend. **d** Visualization of results from GSEA using heatmap. Left (Normalized enrichment score < 0) represents enriched terms in control and right (Normalized enrichment score > 0) represents enriched terms in VPS13B KO organoids.

### Neural spikes and LFP of control and VPS13B KO organoids

Cerebral organoids generate neurons and achieve synaptic maturation, and have previously demonstrated electrophysiological functionality.^52–54^ To evaluate the differences in neural activity between control and VPS13B KO organoids, we recorded spontaneous neural activity. We used a MEMS (Microelectromechanical System)-based silicon neural probe with a 16-channel black Pt microelectrode array, which was inserted into each genotype of 5-month-old organoid. The raster plots indicate that VPS13B KO organoids exhibit significantly higher firing rates than control organoids. Additionally, the amplitude of single spikes recorded from VPS13B KO organoids was higher than that of neural spikes from control organoids (Fig. 6a, b). Furthermore, we measured local field potentials (LFPs) in both control and VPS13B KO organoids. Consistent with the trend of higher firing rates in VPS13B KO organoids, the LFPs also showed higher neural activity with increased amplitudes throughout the organoid structure, from the surface to the center (Fig. 6c). For a more in-depth comparison of LFPs between both groups, we analyzed the LFPs by frequency as well as amplitude. The LFPs from both control and VPS13B KO organoids showed higher magnitudes in the frequency range between 0 and 20 Hz. However, the magnitude of the signal from VPS13B KO organoids was significantly higher than that of control organoids (Fig. 6d-g). We also analyzed the power spectrum of LFPs across different frequency bands. The LFPs from VPS13B KO organoids displayed higher power in the delta band (1-4 Hz), theta band (4-8 Hz), and beta band (12-30 Hz), while power in the alpha band (8-12 Hz) was similar between both types of organoids (Supplementary Fig. 4 and Supplementary Fig. 5). Finally, we compared the neural activity of control and VPS13B KO organoids based on firing rate (Hz), mean spike rate (Hz), mean burst rate (Hz), and burst activity (%). The neural signals from VPS13B KO organoids demonstrated higher burst rates as well as firing rates (Fig. 6h-k). Consistent with electrophysiological observations indicating that VPS13B KO organoids were more excitable, we also observed a reduced proportion of Calbindin-stained inhibitory interneurons in VPS13B KO organoids compared to controls (Fig. 6l, m). These findings collectively suggest that VPS13B KO organoids exhibit hyperexcitability, indicating an alteration in the balance between excitatory and inhibitory neurons.

**Fig. 6.**
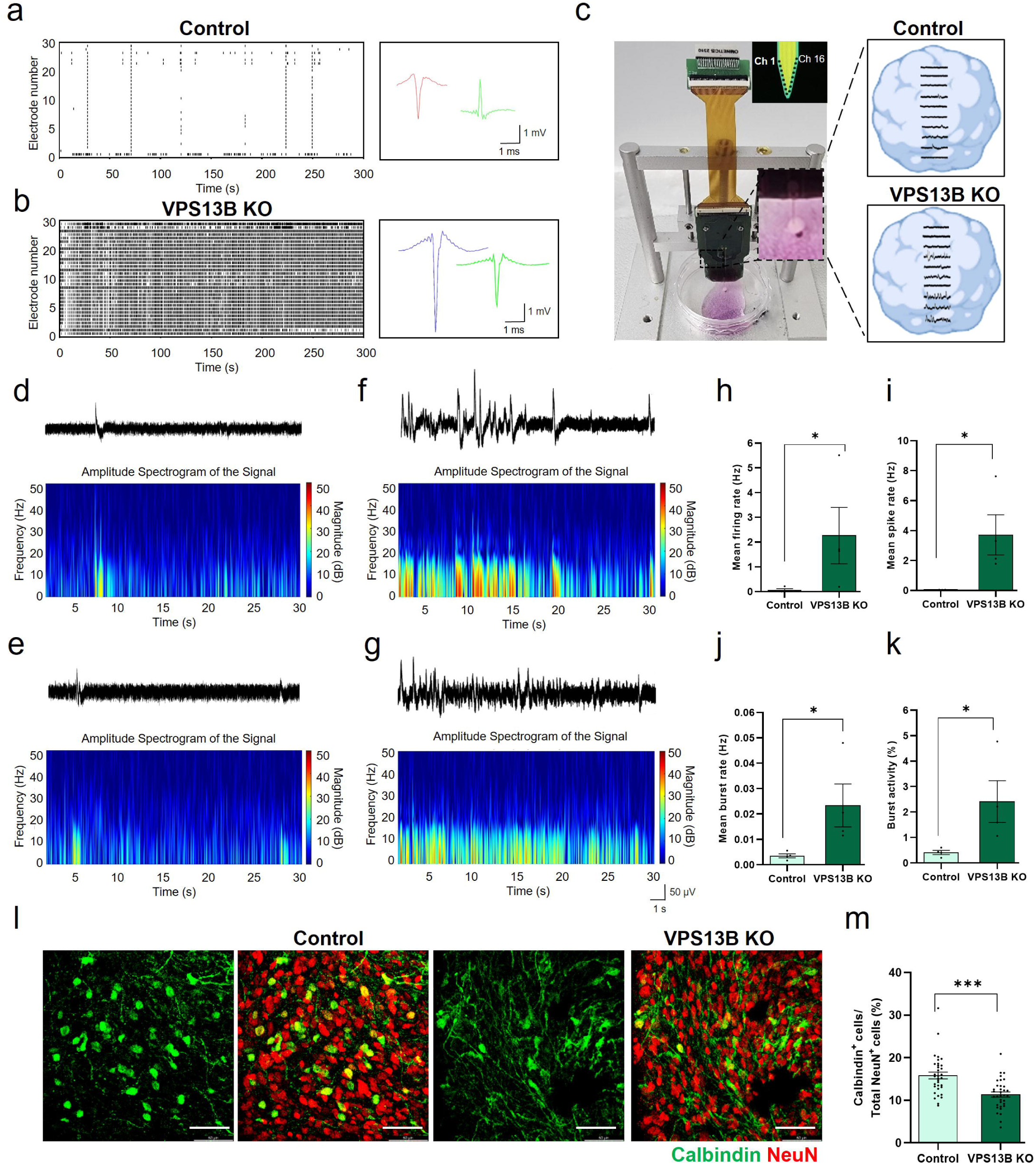
Electrophysiological analysis of control and VPS13B KO organoids. **a** The representative raster plot and neural signals recorded in the 5-month-old organoids. Data were obtained from independent trials (n=5 for control). **b** The representative raster plot and neural signals recorded in the 5-month-old VPS13B KO organoids. Data were obtained from independent trials (n=5 for control and n=5 for VPS13B KO). **c** Photograph of the extracellular recording MEA system using a neural probe integrated with a customized microdrive. A schematic illustration of the neural probe shank with 16-platinum (Pt) microelectrodes. The representative raw local field potential signals (LFP) of the 5-month-old control and VPS13B KO organoids. Data were obtained from independent trials (n=5 for control and n=5 for VPS13B KO). **d, e** The representative raw LFP signals and color-mapped LFP signal amplitude spectrogram of the 5-month-old control organoids. **f, g** The representative raw LFP signals and color-mapped LFP signal amplitude spectrogram of the 5-month-old VPS13B KO organoids. Data were obtained from independent trials (n=5 for control and n=5 for VPS13B KO). **h-k** Bar graphs showing the differences in activities of the 5-month-old control and VPS13B KO organoids (n = 5 independent samples of each controls and VPS13B KO organoids): mean firing rate (h; P = 0.057 between control and VPS13B KO organoids), mean spike rate (I; *P = 0.0342 between control and VPS13B KO), mean burst rate (J; P = 0.0583 between control and VPS13B KO), burst activity (K; P = 0.0519 between control and VPS13B KO). Data are presented as mean ± SD. Statistical significance was tested using Unpaired Student’s t-test for comparing two groups. **l** The reduced number of inhibitory neurons in VPS13B KO organoids was visualized with Calbindin (green) and NeuN (red) staining at 5-month stage (scale bar: 50 μm). **m** Quantification of Calbindin^+^ cells out of total NeuN^+^ cells in control and VPS13B KO organoids (two-tailed unpaired *t*-test; \**p*□<0.001; n=5; number of examined fields = 35 in each group).

## DISCUSSION

Cohen syndrome is one of the rare genetic etiologies of neurodevelopmental disorders (NDDs).^7^ The molecular and cellular pathways driving carrier phenotypes remain poorly understood. VPS13B KO mouse models have traditionally been used to study this syndrome and investigate its disease mechanisms. However, VPS13B KO mice do not faithfully recapitulate human phenotypes: *Vps13b*-mutant mice exhibit microcephaly,^9^ whereas other groups failed to report microcephaly in *Vps13b*^2−/−^ mice.^8,10^ Therefore, we hypothesized that using hPSC-derived neural organoids could provide a more informative model for recapitulating this human disorder. Here, we established a modified protocol for the generation of hANOs to model microcephaly associated with human CS in a tissue-relevant context. By applying this protocol, coupled with genetic manipulation we observed smaller neural organoids in VPS13B KO CRISPR-edited organoids. We identified alterations in neuronal cell growth, including reduction in soma size and impaired neuronal development, as a previously undescribed mechanism contributing to secondary microcephaly. Additionally, electrophysiological analysis revealed hyperexcitability in the VPS13B KO organoids. These results successfully demonstrate the suitability of the modified protocol for disease modelling of CS.

Methods for producing brain organoids have been developed in various ways, enabling the construction of a layered structure similar to the human brain, including processes such as ventricle formation.^55^ Interestingly, the anterior spheroids displayed a short, isolated follicle-like morphology characteristic of ventricular structures, while the posterior spheroids exhibited elongated and interconnected morphologies resembling NT structures. The anterior spheroids subsequently differentiate and mature into hANOs. Following five months of culture, hANOs revealed a significant increase in cortical cell diversity. Moreover, the transcriptomic profile of the 5-month organoid resembles that of the fetal brain during the second trimester of development. The hANOs were larger in size compared to hSCOs (data not shown), suggesting inherent differences in their growth and developmental mechanisms. Analyzing the ventricles of hSCOs and hANOs at advanced maturation stages may yield significant insights into their proliferation dynamics in the future. Our methodology differs from most existing approaches in at least two significant ways. First, we start with initial differentiation in a 2D culture system rather than using 3D embryoid body formation. Then, a specific number of neural-induced cells are re-aggregated to generate uniformly sized organoids. Secondly, our protocol eliminates the need for matrigel embedding, and the hPSCs aggregates are not exposed to guidance cues. Our hANOs generation protocol is robust, enabling the generation of large quantities of organoids per batch that exhibit consistent morphological characteristics.

Microcephaly is one of the core phenotypes in CS patients along with growth delay, hypotonia, and altered memory.^7^ Alterations in progenitor proliferation, cell growth, or differentiation during early development can shape the cortical surface area by modifying cell number, soma size, or morphology.^56^ In our studies, neural progenitor proliferation did not appear to be affected in VPS13B KO organoids. However, it has been reported that neurospheres generated from CS patient-derived iPSCs show a marked reduction in size, along with impaired proliferation of neural progenitor cells.^50^ Given that secondary microcephaly is generally not associated with stem cell proliferation, this discrepancy may highlight the importance of organoid modelling for faithful recapitulation the disease phenotype. Our findings of reduced axon length strongly align with prior reports demonstrating that VPS13B knockdown significantly impairs neurite outgrowth in rat neurons.^3^ The smaller soma size and reduced axon length found in VPS13B KO organoids could be a major contributor to the microcephaly phenotype.

The downregulated genes in VPS13B KO organoids, which are enriched in Gene Ontology terms like chaperone cofactor-dependent protein folding, align with the role of VPS13B in protein folding and transport within the Golgi complex.^57^ The proper trafficking of key receptors and membrane proteins is crucial for neuronal maturation and excitation. Therefore, diminished VPS13B expression and defects in protein processing may hinder the localization or degradation of these essential proteins and receptors, impacting neuronal excitability. Given that approximately 6% of individuals with CS exhibit seizure, ^58–60^ hyperexcitability in VPS13B KO organoids appear to sensitively capture and represent these clinical features in CS patients. Our hANOs protocol enables the production of inhibitory neurons, and the observed hyperexcitability in VPS13B KO organoids seems to be linked to a decreased number of inhibitory neurons. Notably, VPS13B exhibits slightly higher expression in excitatory neurons compared to inhibitory neurons in the adult mouse cortex.^9^ Therefore, it is worth investigating whether VPS13B knockout can differentially affects inhibitory and excitatory neurons, as this may contribute to the imbalance between excitatory and inhibitory signals. Given the lack of available experimental models for studying CS-associated epilepsy, our hANOs model provides a valuable platform for exploring the underlying mechanisms and identifying novel therapeutic targets for this condition. We also propose that the hANOs protocol is sensitive to variations in genetic factors that affect excitatory and inhibitory neurogenesis. Consequently, this culture method holds promise for use in modeling a variety of genetic disorders caused by disruptions in the excitatory-inhibitory balance, as well as for screening potential therapeutic drugs.

## ACKNOWLEDGEMENTS

We are grateful to all laboratory members for their advice and constructive critiques related to this study.

## SUPPLEMENTARY INFORMATION

Supplementary information is available at bioRxiv online.

## DATA AVAILABILITY

The data that support the findings of this study are available upon reasonable requests from the corresponding author.

---

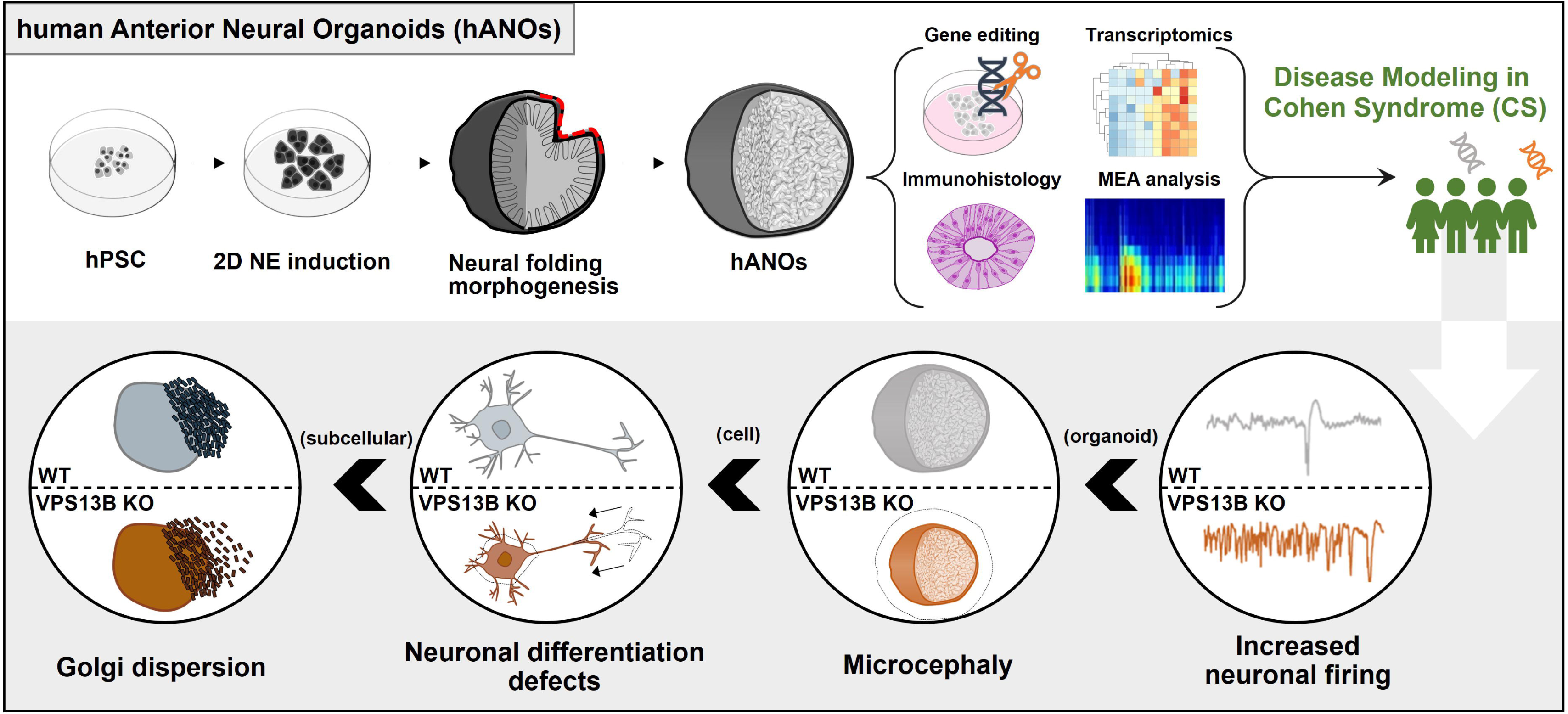

