## supplemental information for "Characterization of Human Anterior Neural Organoids as a Model for Investigating Cohen Syndrome"

**This file includes:**

**Supplementary Fig. 1 to 5**

**Supplementary Fig. 1**

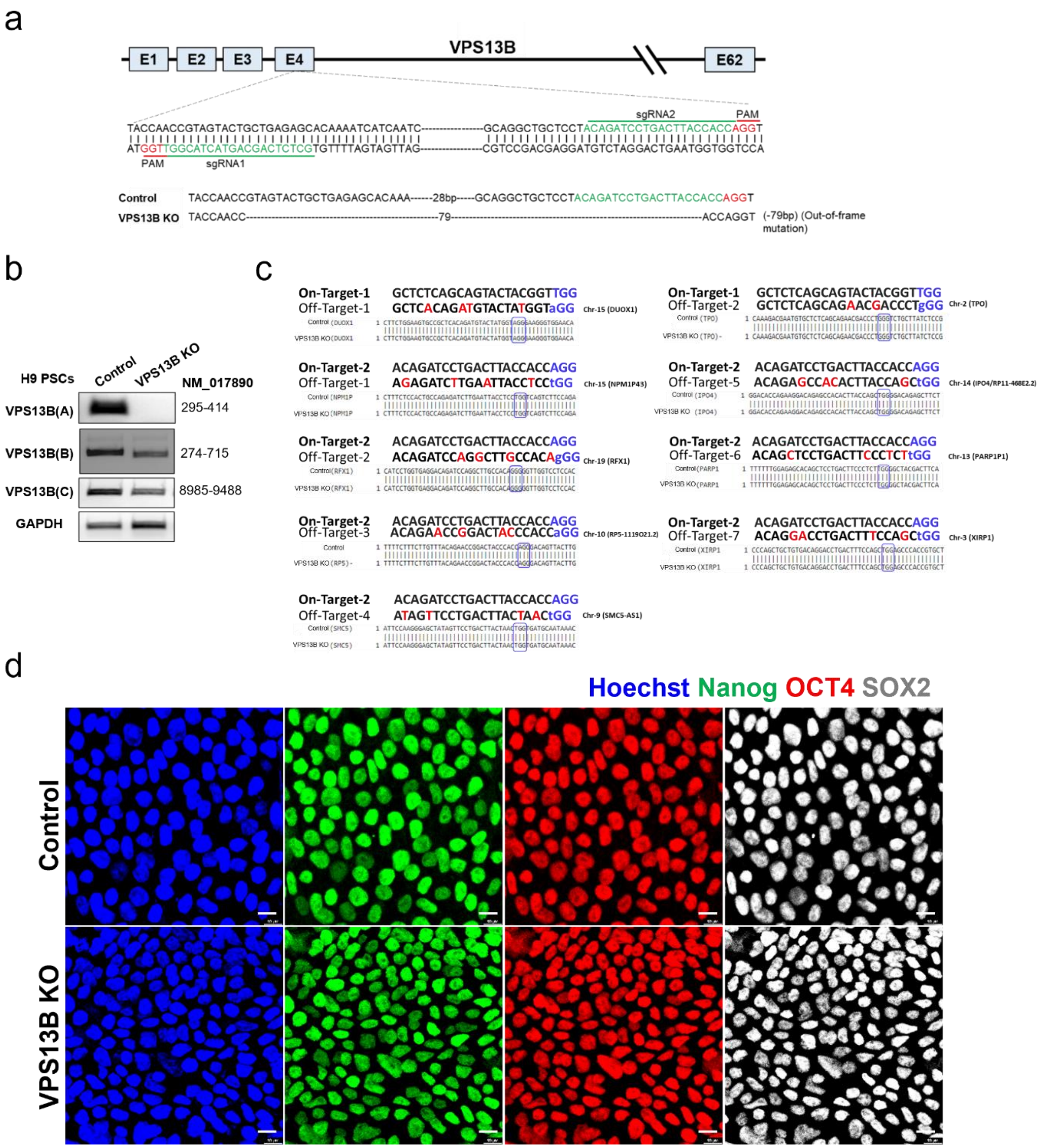


**Supplementary Fig. 1. Generation and validation of VPS13B KO hPSCs. a** The dual gRNAs targeted the exon 4. VPS13B KO mutations occurred as a 79-bp deletion in exon 4 , all of which resulted in a frameshift and led to premature stop codon generation. **b** PCR confirmed the absence of transcripts surrounding the gRNA target region in VPS13B KO hPSCs (panel A). Primers amplifying the C-terminal regions showed the presence of residual transcripts (panel B and C).

These results indicate the C-terminal products are due to alternative splicing process. **c** The sequencing for the off-target effects in the candidate genes indicated the absence of undesired mutations. **d** VPS13B KO hPSCs were generally indistinguishable in expression of pluripotency markers. The VPS13B KO hPSCs were stained with pluripotency markers Nanog (green), OCT4 (red), SOX2(white); scale bar, 10μm.

**Supplementary Fig. 2**

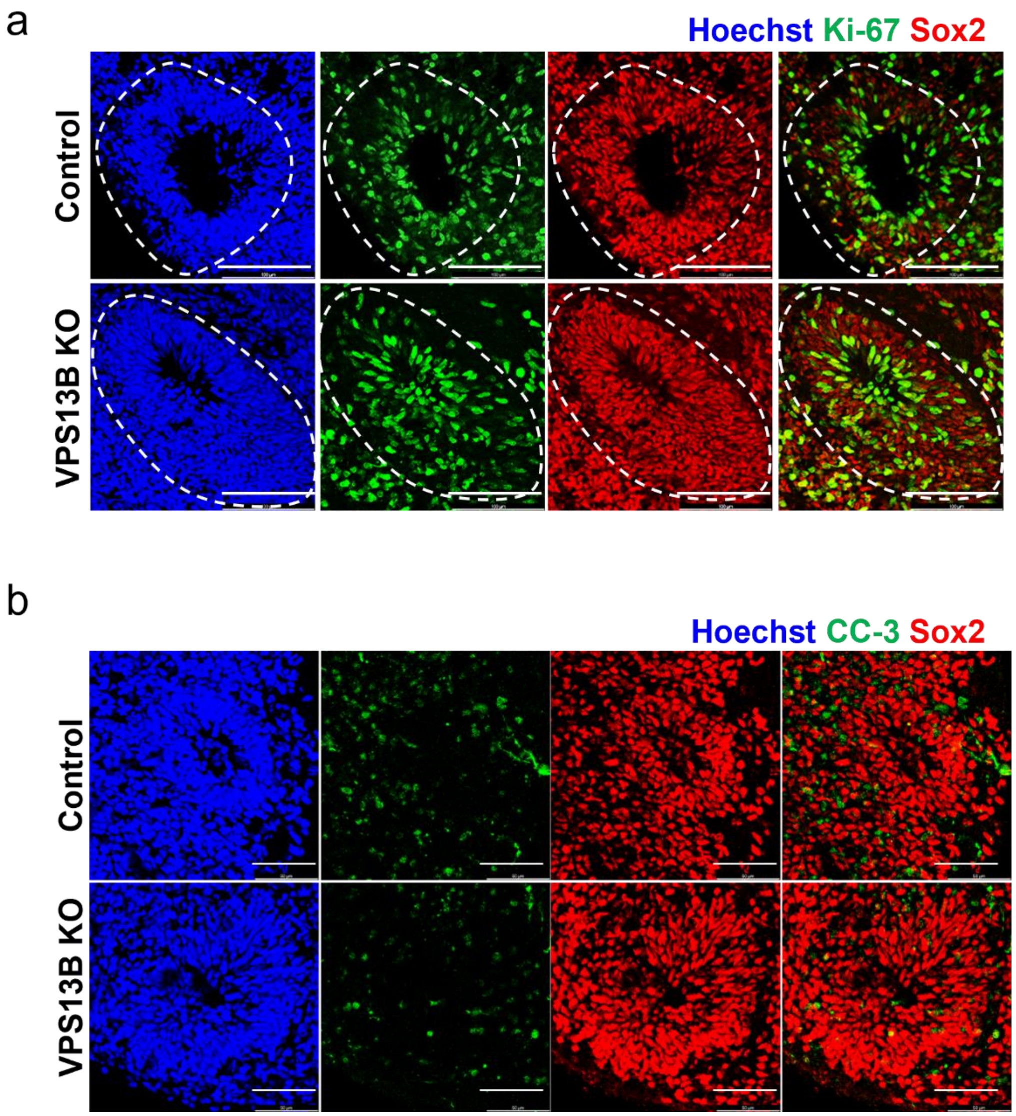


**Supplementary Fig. 2. VPS13B KO does not affect the NPCs proliferation and apoptosis. a** Immunostaining for the NPCs marker Sox2, the proliferation marker Ki67 regions marked with white dotted lines represent VZ like regions; scale bar, 100μm. **b** the apoptosis marker cleaved caspase-3 (CC-3) in 1-month-old control and VPS13B KO organoids. scale bar, 50μm.

**Supplementary Fig. 3**

a

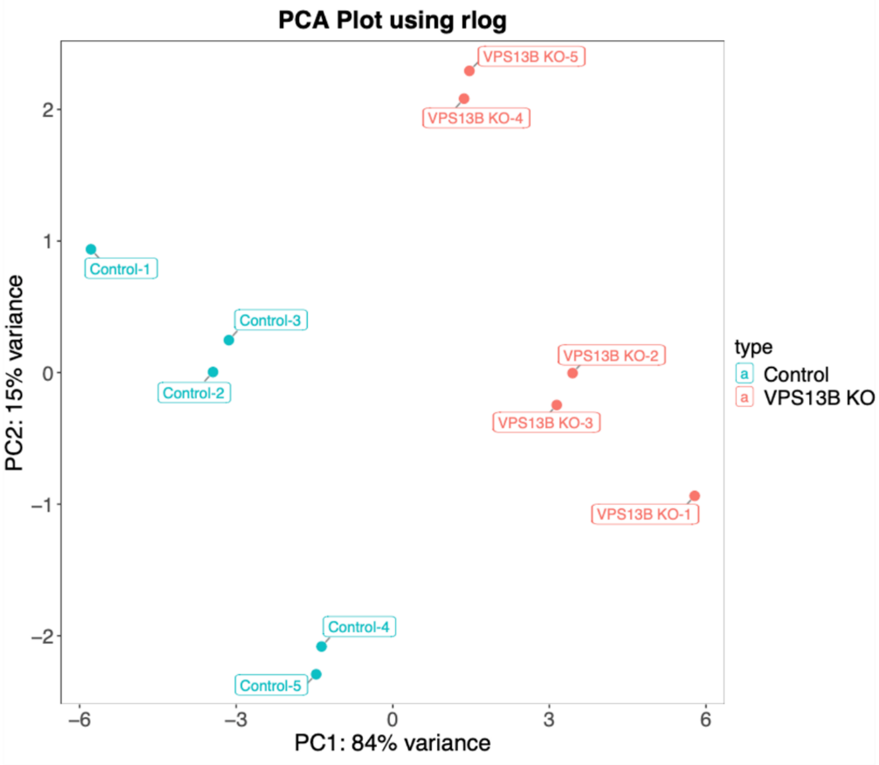


b

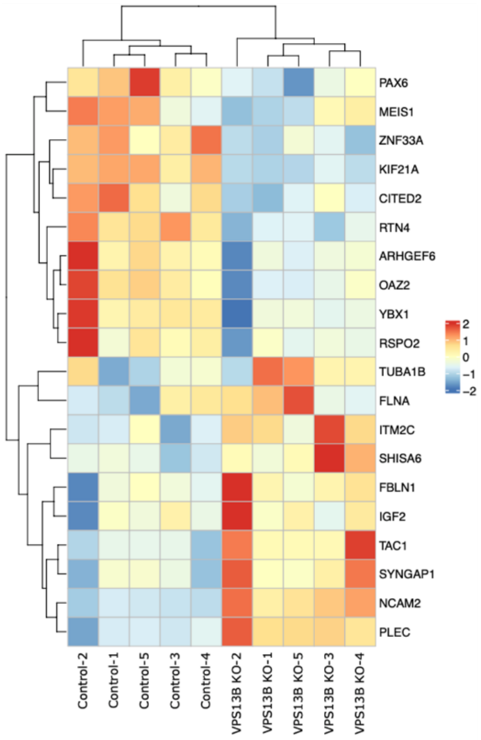

**Supplementary Fig. 3. Principal component analysis (PCA) of control and VPS13B KO organoids. a** PCA plot of RNA-seq data from VPS13B KO and control organoids after batch correction. PC1 and PC2 explain 84% and 15% of total variance, respectively. **b** Heatmap visualization of the top 20 differentially expressed genes (DEGs) distinguishing control and VPS13B KO organoids. Gene expression intensities are displayed as colors ranging from blue for the lowest expression level to red for the highest expression level, as shown in the legend.

**Supplementary Fig. 4**

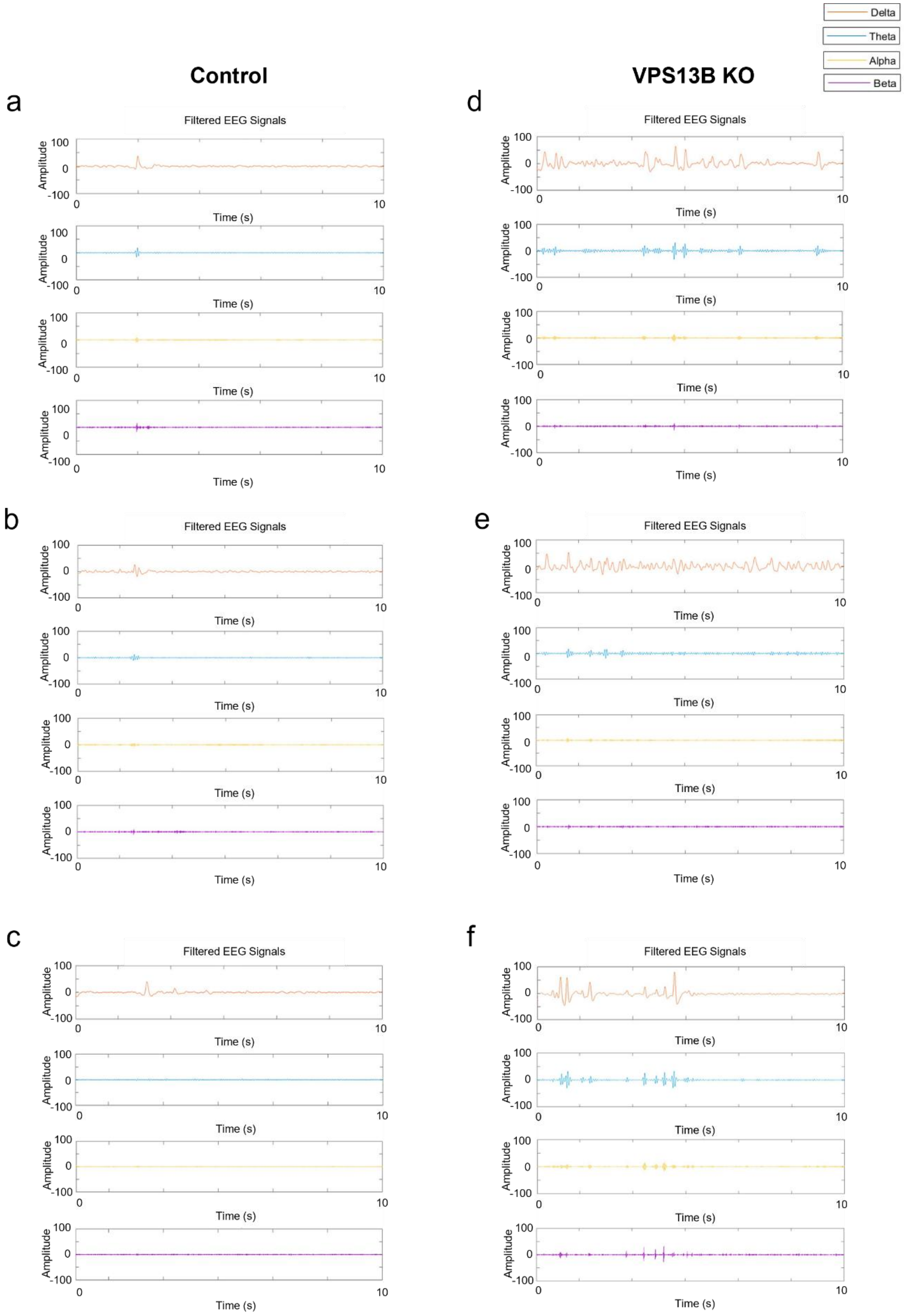

**Supplementary Fig. 4. Filtered LFP signals of control and VPS13B KO organoids in frequency bands (delta (orange): 1-4 Hz, theta (blue): 4-8 Hz, alpha (yellow): 8-12 Hz, beta (purple): 12-30 Hz). a-c** The representative filtered LFP signals recorded in the 5-month-old control organoids. Data were obtained from independent trials ((control (n=5) and VPS13B KO (n=5) organoids)). **d-f** The representative filtered LFP signals recorded in the 5-month-old VPS13B KO organoids. Data were obtained from independent trials (control (n=5) and VPS13B KO (n=5)).

**Supplementary Fig. 5**

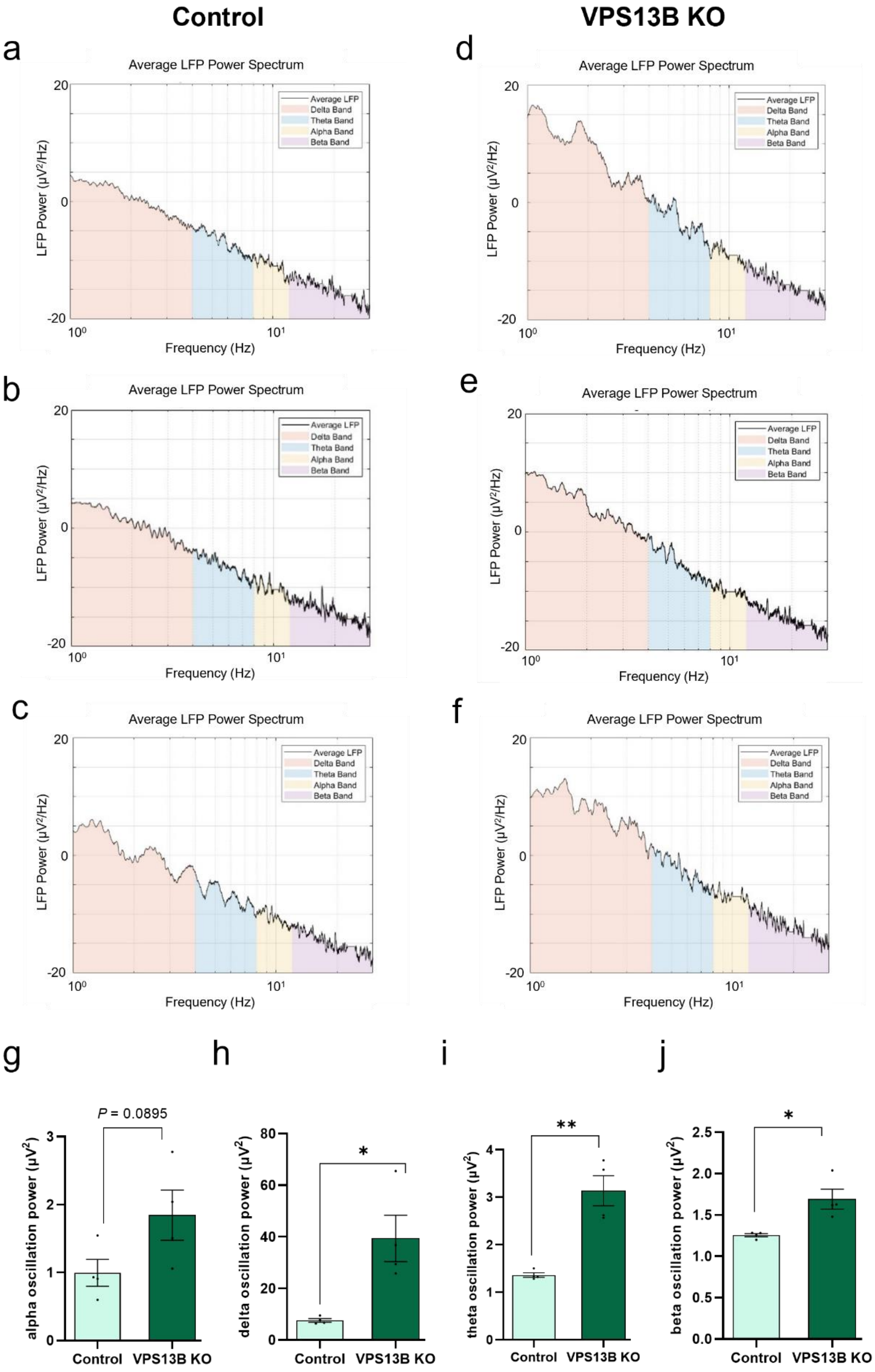

**Supplementary Fig. 5. Electrophysiological analysis of control and VPS13B KO organoids in LFP power spectrum and oscillation powers. a-c** The representative LFP power spectrum of the 5-month-old control organoids. Data were obtained from independent trials (n=5 for control). **d-f** The representative filtered LFP signals recorded in the 5-month-old VPS13B KO organoids. Data were obtained from independent trials (n=5 for CO and n=5 for VPS13B KO). **g-j** Bar graphs displaying the differences in oscillation power between the 5-month-old control and VPS13B KO organoids (n = 5 independent samples of each controls and VPS13B KO organoids): alpha (g; P = 0.0895 between control and VPS13B KO), delta (h; *P = 0.0125 between control and VPS13B KO), theta (I; **P = 0.0014 between control and VPS13B KO), beta (j; *P = 0.0176 between control and VPS13B KO).
